# ERK builds a population of short-lived nascent adhesions that produce persistent edge protrusion and cell migration

**DOI:** 10.1101/2025.11.17.688923

**Authors:** Andrew P. Shepherd, Keith R. Carney, Andrew Elliott, Sangyoon J. Han, Michelle C. Mendoza

## Abstract

Cell migration is realized through the fast and persistent protrusion of a leading edge in the direction of movement. The actin and adhesion structures that build edge protrusions are integrated such that pro-migration signaling pathways must control both assemblies to induce protrusion. Understanding the contribution of adhesion regulation has been complicated by the inability to selectively assay the nascent adhesions that promote edge protrusion. Here, we dissect how the core RAS→RAF→MEK→ERK pathway’s control of nascent adhesions contributes to edge protrusion and cell migration by targeting an ERK FRET biosensor to adhesions and quantifying ERK’s spatial and temporal activity. We find that ERK is activated in the assembling, membrane-proximal region of nascent adhesions through adhesion scaffold paxillin, which interacts with the ERK activator MEK. Tracking nascent adhesion dynamics during cell migration showed that ERK promotes both nascent adhesion assembly and disassembly to create a population of nascent adhesions with short lifetimes. MEK inhibition is partially complemented by expression of a talin R8vvv mutant that increases the nascent adhesion population, demonstrating the significance of ERK’s adhesion regulation for edge protrusion and migration persistence. These findings suggest that when new adhesions initiate, the ERK activation level dictates adhesion assembly and disassembly rates to specifically build nascent adhesions that rapidly turnover, an adhesion population that promotes protrusion persistence and migration.

**Significance:** Cell migration is essential for development, healing, and cancer spread. To move, cells need to build and break tiny structures called nascent adhesions, specifically at their edge that protrudes in the direction of movement. We discovered that ERK, a key signaling protein activated during development and cancer, helps create a fast-turning-over population of these adhesions, which supports steady movement. Using new biosensors and adhesion tracking, we showed ERK is active in the assembling part of the adhesion, where it promotes both assembly and disassembly. Increasing the nascent adhesion population helps cells move when ERK is blocked. This work reveals how ERK controls cell movement by balancing adhesion dynamics and introduces tools to study similar processes in other signaling pathways.

## Introduction

Cell migration is a fundamental biological process essential to embryogenesis and wound healing and adopted by cancer cells during progression to metastasis. Physiological and pathological migration signals induce directionally persistent movement by converting rapid cell edge fluctuations into sustained, or persistent, protrusion in the direction of movement (1, 2). Narrow edge protrusions that occur, split, and retract in multiple directions are converted into persistent protrusion in the forward direction of migration (1, 2).

Signaling pathways control protrusion speed and persistence, and therefore migration, by regulating the underlying actin and adhesion dynamics (1–3). Edge protrusion is driven by actin polymerization against the membrane, which creates a structure called the lamellipodium and pushes the membrane outward (1, 4). At the same time, new adhesions form along the protruding edge and connect the actin network to the substrate (5, 6). Tension in the membrane causes the actin network to undergo retrograde flow and traction force is transmitted across the adhesions as they anchor the actin (5–7). Actin assembly into branched actin structures results in more persistent movement and individual filament elongation drives speed (1, 4, 8, 9). Adhesion-mediated control begins with integrin binding to the extracellular matrix, which induces an active integrin conformation and the recruitment of the force-transducing, actin-binding proteins talin and vinculin, the adaptor protein paxillin, and signaling protein focal adhesion kinase (FAK) (5, 6). This leads to the formation of small, diffraction-limited adhesion complexes (∼120 nm), called nascent adhesions (10–12). As the edge advances, new proteins are recruited into the membrane-proximal “assembling” region (13, 14). These small and short-lived adhesions generate low traction force and are positively associated with protrusion velocity (8, 11, 15–17). Computational modeling predicts that increasing nascent adhesion density, which occurs with increased lifetime, can limit protrusion persistence by tethering actin filaments tightly near the membrane and limiting the ability of new actin monomers to insert between the filament tip and membrane (18).

Nascent adhesions either rapidly turn over (disassemble), move with the leading edge, or mature into large focal adhesions greater than 500 nm in diameter (11, 16). When recruited sequentially, the called nascent adhesions are more likely to turn over than mature into larger focal adhesions (16). Maturation typically occurs at the base of the lamellipodium, where focal adhesions anchor bundled actin fibers, populated with myosin II, to the cell substrate (19–21). Myosin II contracts the actin filaments, causing the assemblies to recruit additional focal adhesion proteins and transmit large traction forces (11, 20). Focal adhesion strength and size predict speed in a biphasic, gaussian relationship, where size increases protrusion speed to a point (11, 20, 22, 23). Actomyosin-mediated stabilization of vinculin is required for stable, polarized protrusion up a chemical concentration gradient or on aligned fibers that direct migration through contact guidance (24, 25). However, excessively large focal adhesions immobilize the actin network, slowing protrusion and inducing retraction (11, 20, 22, 23). Thus, pro-migratory signals must generate a population of nascent adhesions that promotes speed without limiting persistence and which also matures into focal adhesions at a rate that promotes directional movement.

Growth factors, neuropeptides, cytokines, mechanical signals, and integrin engagement with the extracellular matrix activate a handful of key pro-migratory signaling pathways that control both actin and adhesion dynamics (3, 26, 27). Growth factor-responsive kinases, such as extracellular regulated kinase (ERK), protein kinase A (PKA), and protein kinase B (PKB)/Akt, promote adhesion disassembly and their inhibition results in an increase in focal adhesion size and number and reduced cell migration (28–32). While phospho-proteomics has identified numerous substrates within the cell migration machinery (30, 33, 34), how the signaling pathways’ regulation of adhesions generates sustained protrusion and migration remains unknown.

Here, we sought to resolve the role of adhesion regulation for ERK, a principal intracellular signal activated downstream of the growth factor activated and oncogenic RAS→RAF→MEK→ ERK pathway (35, 36). ERK is activated by MEK phosphorylation of the ERK activation loop, which is aided by the joint recruitment of MEK and ERK to scaffolding proteins in different subcellular locations, including adhesions (37–40). Activated ERK is present in lamellipodia protrusions and is required for protrusion velocity and period duration (41–44). ERK promotes branched actin filament assembly through phosphorylation of the WAVE regulatory complex (WRC), which contributes to the velocity and persistence (41, 42). ERK also promotes focal adhesion turnover, as ERK inhibition slows adhesion disassembly and results in larger focal adhesions (32, 41, 45). Whether ERK specifically controls the nascent adhesion population and how this contributes to ERK’s control of protrusion speed and persistence for cell migration is unknown. We addressed these questions by resolving ERK’s spatial and temporal activity in nascent adhesions during active lamellipodia protrusion.

## Results

### Active ERK localizes to the assembling region of nascent adhesions through paxillin

In order to test if active ERK is present in newly forming nascent adhesions, we localized a fluorescence resonance energy transfer (FRET)-based ERK biosensor, called ERK Activity Reporter (EKAR), to adhesions. We fused the Focal Adhesion Targeting (FAT) domain of focal adhesion kinase (FAK) to the N-terminus of EKAREV-NES (EKAR-NES) (46), generating EKAR-FAT (Fig. S1A). The FAK FAT domain can bind talin and paxillin (PXN) in adhesions (47–50). We verified that EKAR-FAT localizes to adhesions by co-transfecting EKAR-FAT and mCherry-talin in COS7 cells and imaging the adhesions by TIRF microscopy (Fig. S1B). Colocalization analysis by Manders overlap coefficient (R^2^=0.847) confirmed specific enrichment of the EKAR-FAT biosensor in talin-labeled adhesions (Fig. S1C). When the analysis was performed with mScarlet-paxillin (Fig. S1D, E), EKAR-FAT and the adhesion marker showed even higher colocalization than with talin, (R^2^=0.9646), suggesting that the EKAR-FAT biosensor is recruited by both talin and paxillin. We verified that EKAR-FAT senses ERK activity similar to the original EKAR-NES by Phos-tag immunoblotting. In serum-starved cells, EGF stimulation increased ERK, EKAR-NES, and EKAR-FAT phosphorylation, while co-treatment with the MEK inhibitor (MEKi, AZD6244) suppressed phosphorylation (Fig. S1F-I). Imaging the FRET from EKAR-NES and EKAR-FAT showed that EKAR-NES sensed ERK activity, as a MEKi-sensitive signal throughout the cell (Fig. S1J). In contrast, EKAR-FAT sensed ERK activity in adhesions (Fig. S1K, yellow arrowheads).

We then used EKAR-FAT to determine if ERK activity is spatially organized within individual adhesions, which might suggest activity associated with adhesion assembly or maturation. We co-expressed EKAR-FAT with mCherry-talin to capture the earliest nascent adhesion assembly state, as paxillin is recruited after vinculin and talin in non-maturing nascent adhesions (16). We then imaged protruding cell edges and quantified EKAR FRET using line scans through individual nascent adhesions, defined as talin puncta <0.5 μm in diameter, newly forming along the direction of protrusion (Fig. 1A) (5, 6, 16). The EKAR acceptor channel reported the biosensor localization and was forward displaced from the peak talin intensity (Fig. 1B). We spatially aligned the intensity peaks of the EKAR channel and FRET ratio (ERK activity) to that of the talin, which revealed that ERK activity was more offset toward the membrane-proximal, assembling region of the nascent adhesion than the sensor itself (Fig. 1B-C). The same localization pattern was observed in nascent adhesions labeled with mScarlet-paxillin (Fig. 1D, E). In this case, the EKAR biosensor’s localization was closely aligned with the peak adhesion signal, consistent with additional EKAR binding to paxillin and recruited sequentially after talin in immature nascent adhesions (Fig. 1F). Nevertheless, the peak EKAR-FAT FRET ratio was offset from paxillin toward the membrane-proximal region, indicating that ERK activity concentrates in the assembling region of new adhesions (Fig. 1G). The finding that peak ERK activity is displaced from biosensor’s localization into the membrane-proximal region of nascent adhesions, irrespective of whether talin or paxillin labeled the adhesions, suggests that ERK activity is enriched in new adhesions following talin and paxillin recruitment (Fig. 1H, I).

**Figure 1:**
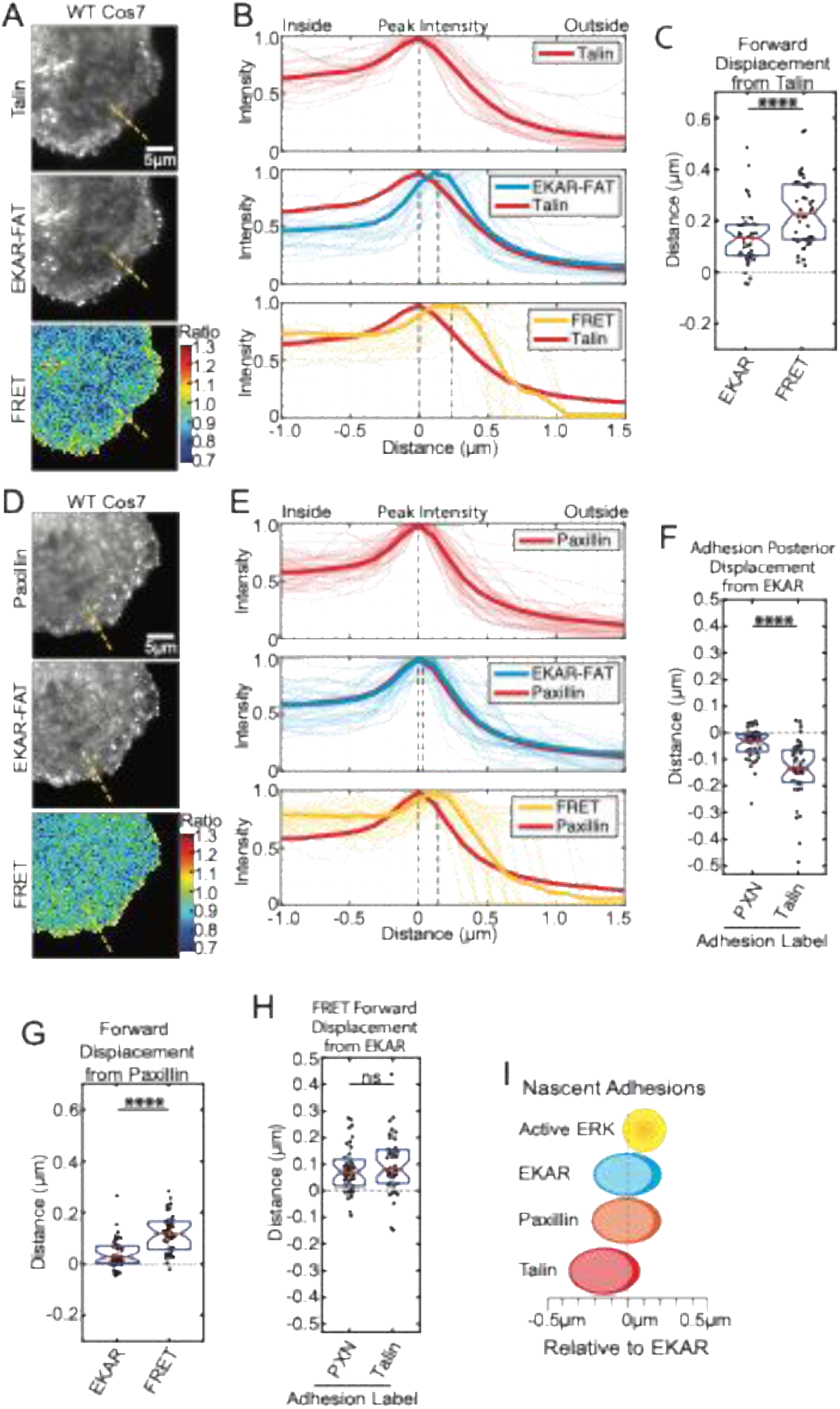
ERK activity localizes to the assembling region of nascent adhesions **(A)** Representative TIRF images of COS7 cells transfected with mCherry-talin and EKAR-FAT and the EKAR-FAT FRET ratio. Scale bar = 5 μm. (**B**) Line scans of assembling nascent adhesions, represented by dashed yellow lines in (A), with peak intensities from the acceptor channel (biosensor localization) and FRET ratio. Peak positions calculated from 3-point polynomial fitting. Bold line is mean. Paired signals are normalized and aligned in space to the mCherry-paxillin peak intensity and position, m=40 adhesions from n=5 cells. **(C)** Forward displacement of peak EKAR-FAT and peak FRET signals. Paired t-test. Boxes are 25^th^ to 75^th^ distribution. Horizontal lines are medians and notches 95% CI around the median. \*\*\*\**p*<0.0001. **(D)** Representative TIRF images (60x) of COS7 cells transfected with mCherry-paxillin and EKAR-FAT and the calculated EKAR-FAT FRET ratio. **(E)** Line scans of assembling nascent adhesions, represented by dashed yellow lines in (A), calculated as in (B), with m=40 adhesions from n=5 cells. **(F)** Quantification of positional displacement of EKAR-FAT relative to mCherry-paxillin and mCherry-talin, from data in (B, E). **(G)** Forward displacement of peak EKAR-FAT and peak FRET signals, calculated as in (C). **(H)** FRET relative to EKAR. Unpaired t-test. (**I**) Model of signal offsets drawn relative to EKAR-FAT.

Given that integrin engagement induces ERK activation and that MEK recruitment to adhesions increases local ERK activity, we hypothesized that ERK is activated within the assembling region of new adhesions during edge protrusion, rather than recruited to the adhesion in an active state. ERK adhesion scaffold proteins paxillin and G-protein coupled receptor kinase interacting protein 1 (GIT1) have been shown to bind to ERK and its activating kinase MEK *in vitro* (51–54) and in the case of Paxillin, promotes ERK activity in adhesions (52). We tested if MEK is also present in the assembling region of newly forming adhesions using bimolecular fluorescence complementation (BiFC) with MEK1 and the paxillin and GIT1 scaffolds. We fused the C-terminal half of the Sapphire fluorescent protein (MEK1-VC155) and fused the N-terminal half of Cerulean or Venus fluorescent proteins to paxillin and GIT1 (PXN-VN173 and GIT1-VN173). We co-expressed the MEK1-VC155 and PXN-VN173 BiFC pair in COS7 cells with mCherry-talin to label adhesions. We then imaged the cells protruding cell edges under TIRF illumination and captured newly forming nascent adhesions. PXN-MEK1 resulted in diffuse cytoplasmic fluorescence and puncta of increased signal intensity at the leading edge (Fig. 2A). We applied the same line scan method and nascent adhesion selection used in the EKAR-FAT study. Line scans drawn perpendicular to the membrane and into the cytoplasm region lacking talin-labeled nascent adhesions showed a baseline BiFC intensity that dropped to the background level outside the cell as the line traversed the edge of the membrane (Fig. 2A, dashed line and 2B, red traces). In contrast, line scans through talin-labelled nascent adhesions showed peaks in BiFC intensity that coincided with talin at the cell edge (Fig. 2A arrowheads, 2B blue traces, and 2C). Similar results were found with the GIT1-VN173 and MEK1-VC155 BiFC pair (Fig. 2E-G), indicating that MEK interacts with both paxillin and GIT1 in nascent adhesions. Furthermore, the paxillin and GIT1 BiFC intensity peaks were similarly forward displaced from talin (Fig. 2G-I), consistent with MEK recruitment to the actively assembling region of nascent adhesions.

**Figure 2:**
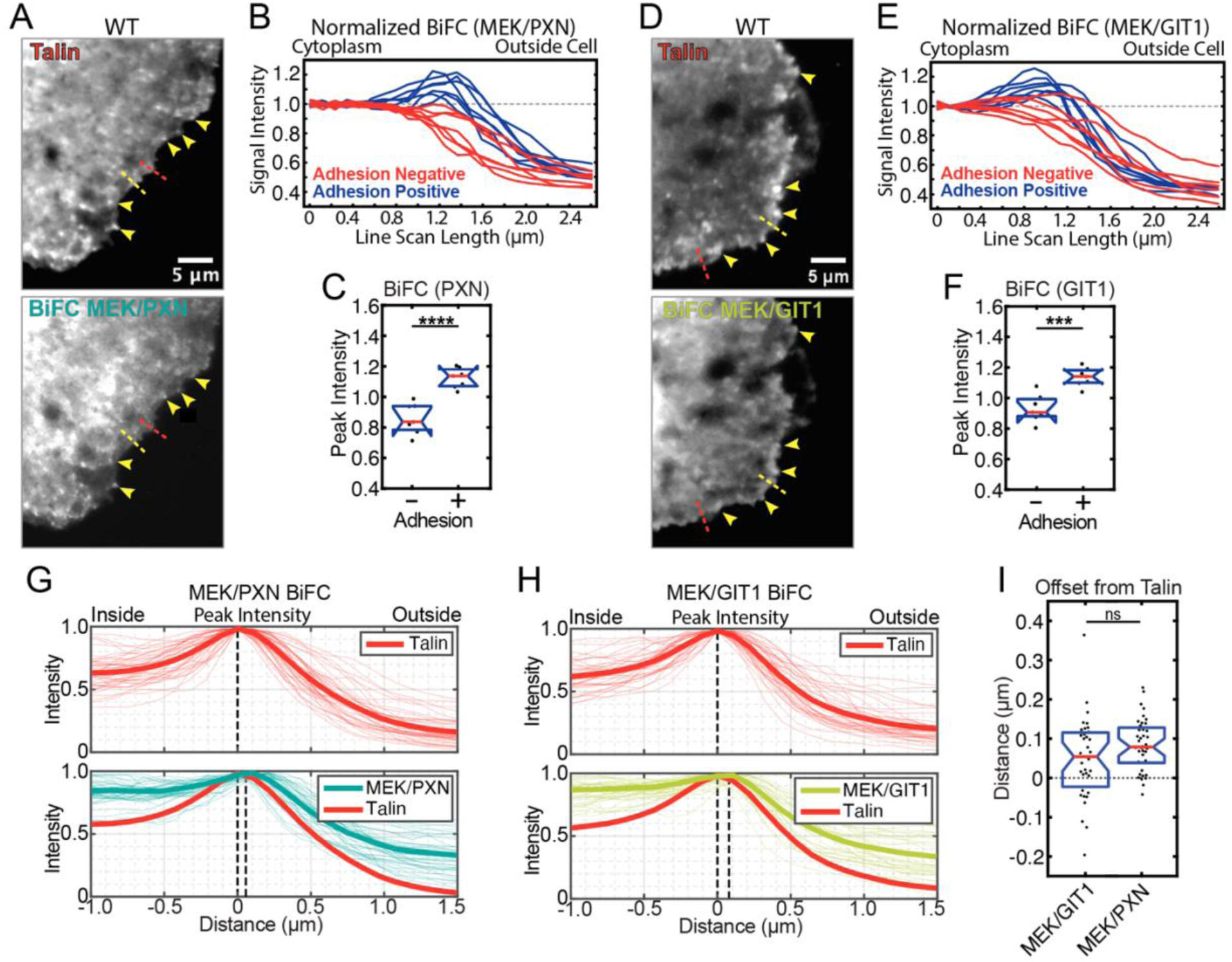
MEK interacts with ERK scaffolds in nascent adhesions COS7 cells were co-transfected with BiFC pairs Sapphire VC155-C1-MEK1 and Cerulean-N173-N1-PXN or Venus-N173-N1-GIT1, and mCherry-talin as a fiducial marker of adhesions. **(A)** Representative TIRF images (60x) of mCherry-talin and MEK/PXN BiFC. Scale bars = 5 μm. **(B)** Line scans of the BiFC signal in (A), normalized and aligned in space to the mCherry-talin line’s peak intensity and position. Adhesion negative lines crossed the cell edge without intersecting a nascent adhesion, represented by red dashed line in (A). Adhesion positive lines crossed the cell edge and intersected an mCherry-talin nascent adhesion, exemplified with yellow dashed line (A). Arrowheads show corresponding features between channels in (A). BiFC signal was normalized to the background intensity of the cell interior, n=40 line scans for both positive and negative, from n=7 cells. **(C)** Mean peak intensity in BiFC signal from lines scans in (B). Boxes represent the 25^th^ to 75^th^ distribution. Horizontal lines are medians. Notches indicate 95% CI around the median. Unpaired t-test. \*\*\*\**p*<0.0001. **(D-F)** Representative images, lines scans, and BiFC intensity values from COS7 cells expression mCherry-talin and MEK/GIT1, analyzed and quantified as in (A-C). \*\*\**p*<0.001. **(G, H)** MEK/PXN and MEK/GIT1 BiFC intensity peaks, from lines scans in (A, D), Peak positions calculated from 3-point polynomial fitting. Bold line is mean. Plots of paired signals of normalized peak intensities from BiFC channels were plotted relative to mCherry-talin, m=40 adhesions from n=5 cells. **(I)** Forward displacement of peak BiFC signals, quantified relative to mCherry-talin. Unpaired t-test.

In order to test whether paxillin and GIT1 are required for ERK activation in newly forming adhesions, we generated COS7 cells lacking PXN and GIT1 using CRISPR/Cas9. For each scaffold, we generated knockout clones using guide RNAs targeting two different exon regions and confirmed loss of paxillin and GIT1 protein expression by Western (Fig. 3A, B). The COS7 paxillin and GIT1 knockouts (PXN KO and GIT1 KO) exhibited reduced migration velocity and directional persistence, compared to WT controls (Fig. S2), phenocopying paxillin and GIT1 loss in other cell types by random walk migration assay (54, 55). Representative PXN clone 1-5 and GIT1 clone 1-4 were selected for further experiments.

**Figure 3:**
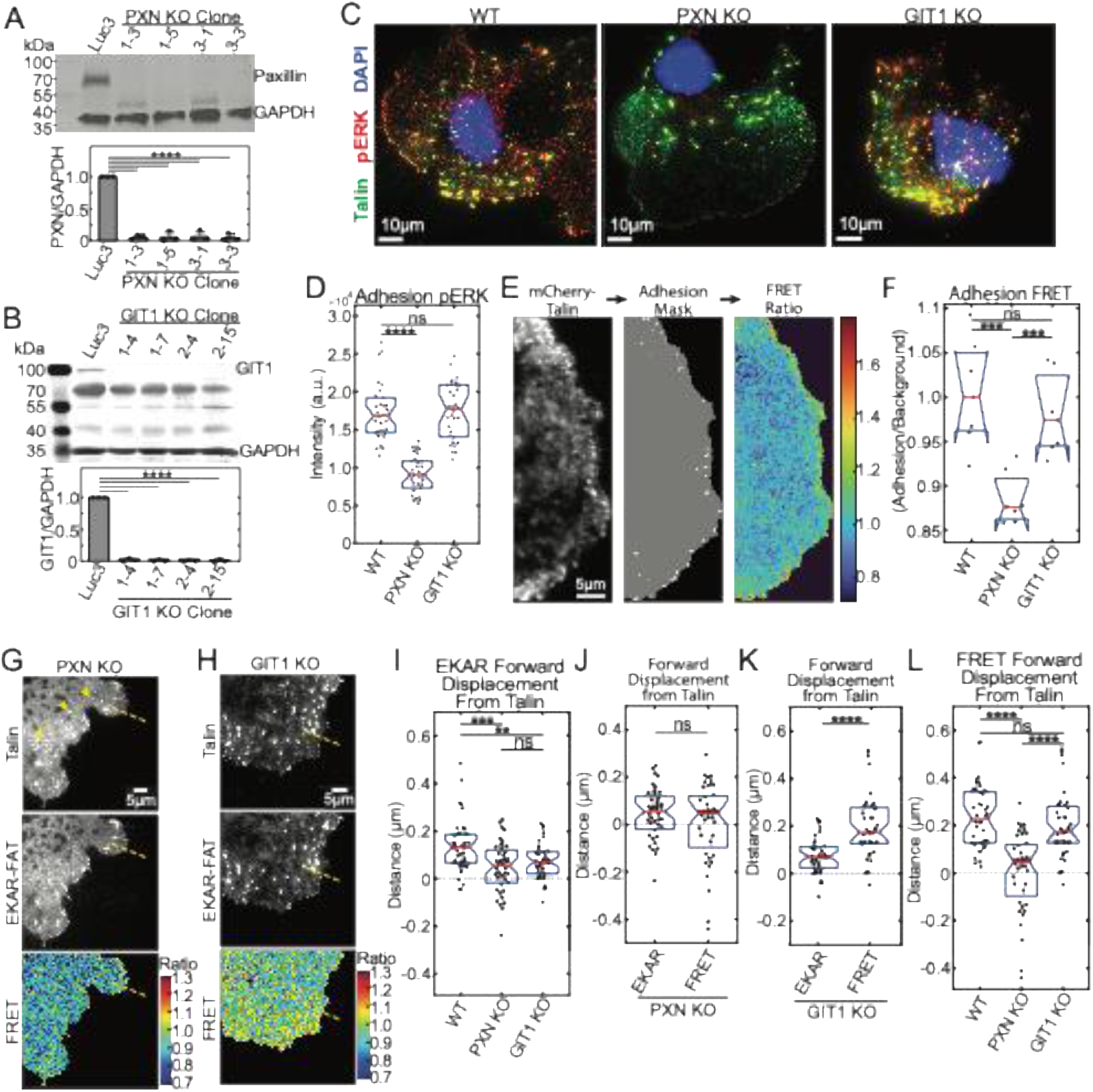
Paxillin scaffolds ERK activity in the nascent adhesions of protruding edges **(A, B)** Representative Western blots and quantification of paxillin and GIT1 expression in COS7 clones transfected with gRNAs targeting *luciferase*, *paxillin*, or *git1*, n=3 experimental replicates. Error bars are SD. **(C)** Representative TIRF immunofluorescence images (60x) of COS7 WT, COS7 PXN KO, and COS7 GIT1 KO cells labelled for pERK (T202/Y204) (red), mCherry-talin (green) and DAPI (blue). **(D)** Mean pERK intensity in adhesions segmented by top hat filter from cells in (C), n=30 cells per condition. Unpaired t-test. **(E)** Adhesion FRET analysis in cells co-transfected with EKAR-FAT and mCherry-talin: protruding cell edge with mCherry-talin labelled adhesion is used to generate adhesion masks and the FRET ratio is calculated in the masked adhesions. Scale bar = 5 μm. **(F)** Adhesion FRET ratio / cytoplasmic FRET, normalized to that of WT COS7 cells. n=7 protrusions from each cell line. Unpaired t-test. (**G, H**) Representative TIRF images (60x) of live COS7 PXN KO and GIT1 KO cells co-transfected with mCherry-talin and EKAR-FAT, with the calculated FRET ratios. Dotted yellow line represents a typical line scan bisecting an EKAR+ assembling adhesion along a protruding edge. Yellow arrowheads highlight interior adhesions labeled by mCherry-talin and not labeled by EKAR-FAT. (**I**) Line scans were normalized to peak intensity and position of mCherry-talin from n=40 adhesions in 5 cells. Peak position calculated from 3-point polynomial fitting. Quantification of EKAR-FAT signal displacement relative to mCherry-talin. One-way ANOVA. (**J, K**) Quantification of signal displacement of EKAR-FAT and calculated FRET signals relative to mCherry-talin in PXN KO and GIT1 KO cells using paired t-test. (**L**) Quantification of combined displacement of the peak FRET signals relative to Talin in 3J and 3K. One-way ANOVA. \*\**p*<0.01, \*\*\**p*<0.001, \*\*\*\**p*<0.0001.

We tested if paxillin and GIT1 are required for the localization of active ERK in adhesions by immunofluorescence for phosphorylated ERK (p-ERK) and talin (Fig. 3C). Quantification of the p-ERK intensity specifically in nascent adhesions within 5 μm of the cell edge revealed that PXN KO cells harbored less ERK activity in nascent adhesions than wildtype COS7 cells, while GIT1 KO cells were indistinguishable from wildtype cells (Fig. 3D). We next tested if paxillin or GIT1 scaffold ERK activation in migrating COS7 cells using the EKAR-FAT biosensor. We transfected COS7 WT, PXN KO, and GIT1 KO cells with EKAR-FAT and mCherry-talin, which we used to identify and mask nascent adhesions along protruding edges. We calculated the FRET, or ERK activity, within adhesions (Fig. 3E). Using background, cytoplasmic FRET to normalize expression levels between cells, we found that ERK activity in PXN KO cells was lower than WT cells, while ERK activity in GIT1 KO cells was indistinguishable from WT cells (Fig. 3F). While both paxillin and GIT1 form complexes with MEK in adhesions, the loss of nascent adhesion-localized p-ERK and loss of EKAR-FAT FRET signal specifically in the PXN KO cells suggests that ERK activation in nascent adhesions is PXN-dependent, but not GIT1-dependent.

We next sought to determine if paxillin specifically mediates ERK activation within the assembling region of nascent adhesions. Using line scans, we analyzed the relative positions of mCherry-talin, EKAR-FAT, and the peak FRET signal for ERK activity in PXN KO and GIT1 KO cells (Fig. 3G, H). In both PXN KO and GIT1 KO cells, the EKAR-FAT biosensor was present in newly assembling adhesions, but its localization was no longer offset toward the leading edge (membrane-proximal) relative to talin (Fig. 3I-K and S3A, B). We noted that in the PXN KO cells, some smaller adhesions at the leading edge, labeled with mCherry-talin failed to recruit EKAR-FAT (Fig. 3G, compare Talin signal in top panel arrowheads to presence of EKAR-FAT in middle panel). This indicated that in the absence of paxillin, the FAT domain of EKAR-FAT variably associates with talin or another focal adhesion protein where it can report out ERK activity. In PXN KO cells, the remaining FRET signal peaked at the same location as the biosensor itself (Fig. 3J and S3A). In GIT1 KO cells, ERK activity remained forward displaced from the biosensor (Fig. 3K and S3B). Similarly, the forward displacement of ERK activity from talin was reduced in the PXN KO cells, but not in GIT1 KO cells (Fig. 3L). Thus, only paxillin is required for ERK activation in the membrane-proximal, assembling region of nascent adhesions.

### ERK increases the nascent adhesion population by promoting both assembly and disassembly

Since active ERK localizes to the assembling region of nascent adhesions, we tested whether ERK activity plays a role in nascent adhesion assembly, turnover, and maturation. We employed a machine learning–based approach from Han et al., 2021 to track adhesion dynamics, categorize the adhesions into eight distinct classes based on changes in size, shape, and intensity over time, and analyze their relationship with protrusion behavior. COS7 cells were transfected with mScarlet-paxillin to label adhesions, adhesions were monitored during cell migration by TIRF timelapse imaging, and cells were treated with DMSO control or MEKi (AZD6244) to assess ERK’s contribution to adhesion dynamics. Regions of interest (ROI)s for analysis were selected from protruding areas at the cell edge (Fig. 4A and S4A). MEKi significantly reduced edge protrusion, reducing mean and maximum protrusion velocities and causing a 23.1% decrease in protrusion persistence compared to control DMSO-treated cells, mirroring our previous study in PtK1 cells (42) (Fig. S4B-E). The protrusions of control, DMSO-treated cells were populated with small, rapidly turning over adhesions (Fig. 4B and Supplemental Movie 1). In contrast, the stalled edges of cells treated with MEKi accumulated larger, stable adhesions (Fig. 4A, B and Supplemental Movie 2). Quantification of the nascent and focal adhesion number showed MEKi-treated cells have a reduced nascent adhesion / focal adhesion ratio compared to DMSO-treated cells, indicating that MEK inhibition causes cells to shift nascent adhesions to stable adhesions (Fig. 4C).

**Figure 4:**
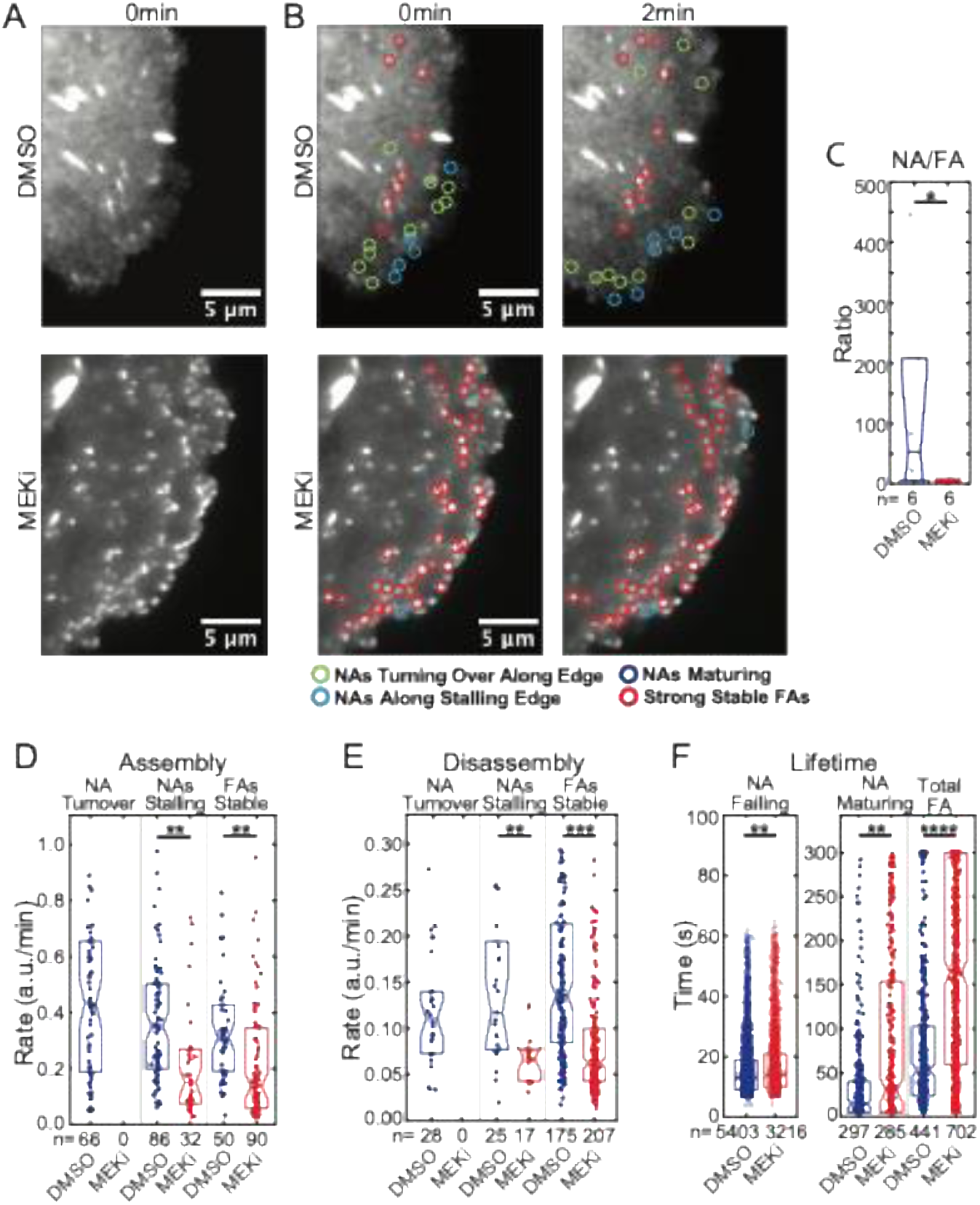
ERK builds a population of nascent adhesions by promoting both assembly and disassembly COS7 cells were transfected with mCherry-paxillin and treated DMSO or MEKi (AZD6244, 5 μM). Timelapses were acquired for 5min at 10s intervals using TIRF microscopy (60x), n=6 cells per condition. Adhesion tracking based on all detected point sources within 7μm of the cell edge. Point source signals exhibiting characteristics of nascent adhesions (NAs) or focal adhesions (FAs) were assigned to representative groups following the classification schema in Han, *et al.*, 2021. **(A)** Representative images of analyzed protrusions and **(B)** visualized adhesion classification. **(C)** Ratio of NAs / FAs in protrusions in each cell. Unpaired t-test. **(D)** Assembly rate and **(E)** disassembly rate for individual adhesions of each class of interest, for each condition. K-S test. **(F)** Adhesion lifetimes. K-S test. Boxes represent the 25^th^ to 75^th^ distribution. Horizontal lines are medians and notches 95% CI around the median. \**p*<0.05, \*\**p*<0.01, \*\*\**p*<0.001.

We then quantified the adhesion assembly and disassembly rates and lifetimes. In the control, DMSO-treated COS7 protrusions, we identified rapidly turning over nascent adhesions, nascent adhesions along stalling edges, and stable focal adhesions (Fig. 4D). MEKi resulted in a loss of rapidly turning over nascent adhesions (Fig. 4D, E). MEKi also reduced both the assembly and disassembly rates of nascent adhesion along stalling edges and focal adhesions, compared to control, DMSO-treated cells (Fig. 4D, E). We analyzed the lifetime of the adhesion population by classifying structures into nascent adhesions that fail or turn over during the imaging acquisition, nascent adhesions that increase in size to be re-classified as focal adhesions during the imaging acquisition, and adhesions that appear large enough to be focal adhesions during the entire timelapse. This revealed that MEKi increased both nascent and focal adhesion lifetime (Fig. 4F). Thus, ERK promotes both the assembly and turnover of nascent adhesions, shifting the total adhesion population to one comprised of more nascent adhesions with shorter lifetimes.

### Increasing the nascent adhesion population partially rescues the protrusion and migration defects caused by MEK inhibition

We hypothesized that ERK’s activity in building a short-lived nascent adhesion population contributes to ERK’s promotion of protrusion speed and duration for effective cell migration. If true, increasing the population of nascent adhesions should complement the suppressed protrusion velocity and persistence caused by MEK inhibition. We tested this hypothesis by using the talin R8vvv mutant, which harbors the T1502V, T1542V, and T1562V mutations that remove the threonine belt needed for talin unfolding and prevent force-independent vinculin binding and maturation of nascent adhesions into focal adhesions (16). We generated COS7 cells with a stable talin1 knockdown (shRNA talin) and then transfected wildtype talin and the R8vvv mutant, tagged with mNeonGreen (mNG), into the shRNA talin cells (Fig. 5A). Talin1 knockdown changed the cell morphology to a condensed shape lacking broad lamellipodia and reduced cell migration (Fig. S5A-D). Reintroduction of talin resulted in a full-length Talin1-mNG and Talin1-R8vvv-mNG expression, as well as a cleavage product for each talin construct, presumably due to the described calpain-mediated cleavage (56). Nevertheless, expression levels remained above endogenous levels and comparable between constructs (Fig. 5A).

**Figure 5:**
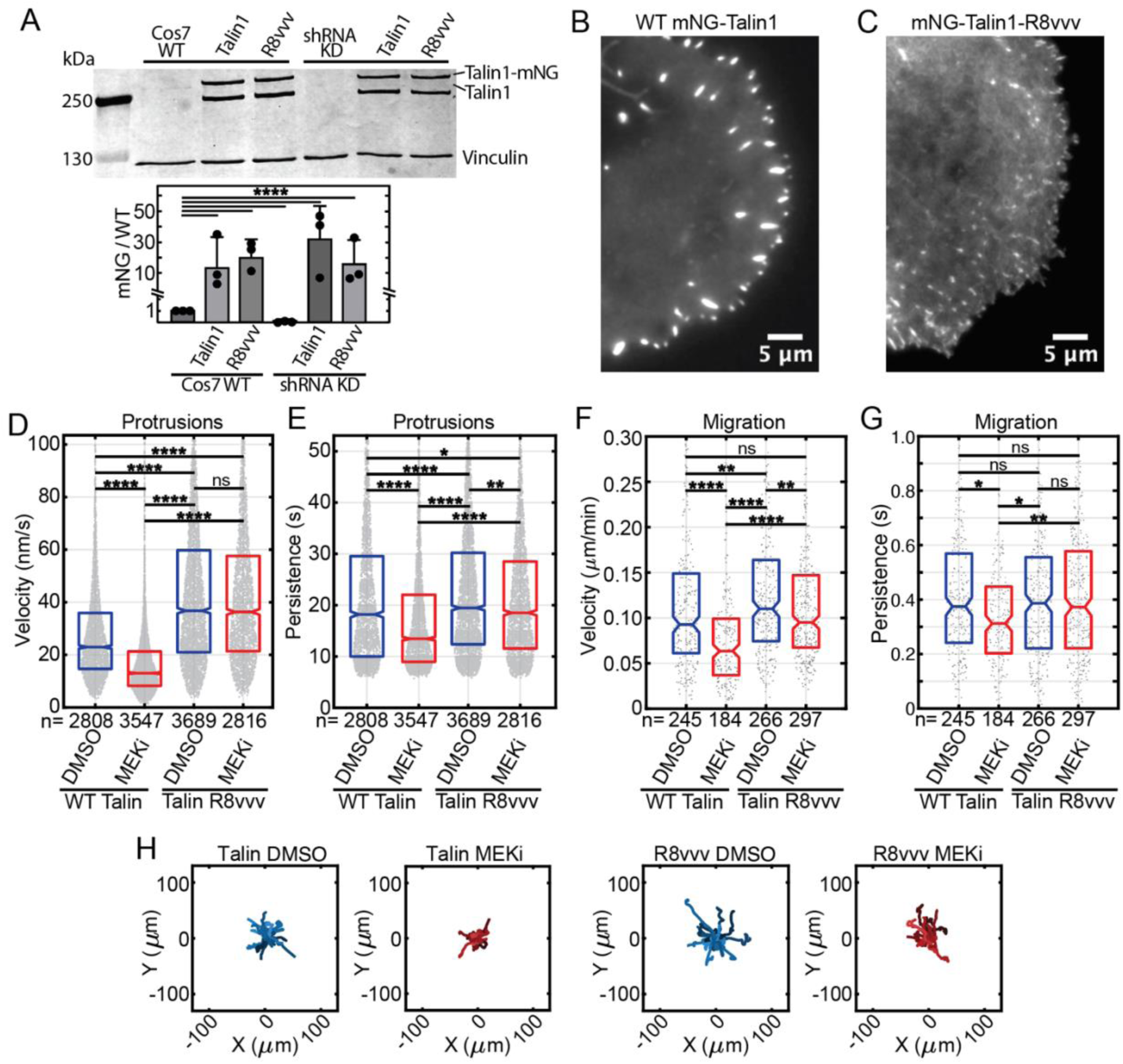
Increasing the population of nascent adhesions compensates for loss of ERK to drive edge protrusion and migration COS7 cells expression Talin1-shRNA were transfected with WT mNG-Talin1 or mNG-Talin1-R8vvv. Representative Western blot and quantification of endogenous talin, talin1 knockdown transient transfection expression of mNG-Talin1 and mNG-Talin1-R8vvv (**A**). Median mNG expression normalized to endogenous talin. Error bars SD. Representative TIRF microscopy images (60x) of adhesions in protrusions of talin1-shRNA cells transfected with mNG-Talin1 (**B**) or mNG-Talin1-R8vvv (**C**). Talin-shRNA KD cells were transfected with mNG-Talin1 or mNG-Talin1-R8vvv 48 hours prior to imaging. Confocal images (60x) were acquired at 5 s intervals for 8 min. Areas of active leading-edge protrusion were cropped for analysis of 8 replicates of each condition and n= numbers of windows tracked. Mean protrusion velocity (**D**) and persistence (**E**). K-S test. Talin-shRNA KD cells were transfected with mNG-Talin1 or mNG-Talin1-R8vvv 48 hours prior to imaging and treated with DMSO or 2 μM MEKi (AZD6244) 30 min prior to imaging. Cells were imaged at 10 min intervals for 12 hours (20x phase contrast and fluorescent widefield) and tracked using MATLAB software. Random walk migration MSD velocity (**F**) and persistence (**G**). K-S test. Number of cells tracked and analyzed (n) shown for each condition. Rose plots display 20 cell tracks with speeds nearest to the median for each condition (**H**). Boxes represent the 25^th^ to 75^th^ distribution. The horizontal line indicates the mean or median, as stated, of each group. Notches indicate 95% CI around the median. \**p*<0.05, \*\**p*<0.01, \*\*\**p*<0.001, \*\*\*\**p*<0.0001.

TIRF imaging of the adhesions confirmed that WT mNG-Talin1 presented as larger, more stable focal adhesions at the cell periphery, while mNG-Talin1-R8vvv appeared in small, punctate adhesions consistent with nascent adhesion structures (Fig. 5B, C), consistent with the R8vvv mutant generating a nascent adhesion-dominant adhesion population. As previously reported, R8vvv-expressing cells exhibited faster protrusion velocity compared to cells expressing WT talin (Fig. 5D, (16)). The R8vvv-expressing cells also exhibited increased protrusion persistence compared to WT talin-expressing cells (Fig. 5E). We then tested if the increased nascent adhesion-shifted population R8vvv-expressing cells could rescue MEK inhibition and the loss of ERK activity. MEK inhibition in WT talin-expressing cells reduced protrusion velocity and persistence (Fig. 5D, E), similar to MEK inhibition in COS7 cells (Fig. S4E). In contrast, R8vvv-expressing cells maintained their protrusion velocity under MEK inhibition and exhibited a 5.0% decrease in protrusion persistence (Fig. 5E). Thus, increasing the population of nascent adhesions preserves protrusion dynamics in the absence of ERK signaling. We tested whether the complementation of MEK-inhibited cells’ nascent adhesion population and protrusion dynamics recovered their migration defects. Consistent with the protrusion data, R8vvv-expressing cells migrated faster and more persistently than WT talin rescue cells (Fig. 5F-H and Supplemental Movies 3 and 4). MEK inhibition reduced migration velocity 28.1% and persistence 16.8% in the WT talin rescue cells (Fig. 5F, G). However, MEKi reduced migration velocity only 15.4% and had no effect on migration persistence in the R8vvv-expressing cells (Fig. 5G, H). Thus, nascent adhesions can support cell migration in the absence of ERK signaling.

## Discussion

We have found that active ERK is present in the assembling region of newly forming nascent adhesions, where it promotes both assembly and disassembly to create a nascent adhesion population that promotes protrusion persistence for cell migration. Previous measurements of ERK-controlled adhesion dynamics relied solely on normalized intensity changes at a fixed position of sliding adhesions and concluded that ERK strictly promotes adhesion disassembly (28, 32, 45). We identified ERK’s role in adhesion assembly by quantifying rate constants on the tracked, moving positions of adhesions in a sub-resolved manner. Thus, ERK builds short-lived nascent adhesions that support stable protrusion. This finding supports the model that pro-migratory signaling programs control both actin and nascent adhesions to support protrusion and migration persistence.

Live-cell imaging of the EKAR-FAT biosensor revealed that ERK activity is spatially enriched in the membrane-proximal, assembling region of adhesions. We determined that paxillin is the key scaffold protein that binds MEK and mediates ERK activation in the assembling region of nascent adhesions. Paxillin knockout, but not GIT1 knockout, reduced ERK activity in nascent adhesions and abrogated active ERK’s localization to the assembling region of nascent adhesions. This suggests that paxillin functions, in part, by enriching ERK activity in the assembling region of nascent adhesions, thereby promoting adhesion turnover and protrusion persistence to increase directional migration. We previously found that when paxillin is recruited concurrently with talin and simultaneously with the initial rise in traction, the nascent adhesions are more likely to mature into focal adhesions (16). Thus, paxillin recruitment without MEK→ERK activation may function similar to co-recruitment with talin, in which traction induces adhesion assembly, reinforcement, and stabilization. We found that GIT1 interacted with MEK in adhesions and was required for edge protrusion and cell migration. However, GIT1 did not control ERK activation in actively protruding cell edges. While GIT1 can bind paxillin, GIT1 appears to be recruited to larger structures called focal contacts, which develop from stabilized nascent adhesions and are in the process of transitioning to focal adhesions (54, 57). Our findings suggest that GIT1’s control of focal contact turnover is ERK-independent.

Our conclusion that ERK promotes the formation and turnover of short-lived nascent adhesions at the leading edge, which contributes to protrusion persistence, was supported by multiple findings. MEK/ERK inhibition abolished the formation of rapidly turning over nascent adhesions, reduced nascent adhesion disassembly rates, and resulted in a shift toward longer-lived, stable focal adhesions. This shift corresponded with diminished protrusion velocity and persistence, as well as less persistent migration. We have previously shown that ERK contributes to lamellipodial actin polymerization as part of its mechanism in driving protrusion persistence (41, 42). Here, we directly tested if ERK’s generation of rapidly turning over nascent adhesions also contributes to its persistent edge protrusion. The effects of ERK inhibition on adhesion turnover and migration were functionally recovered by increasing the population of small, rapidly turning over adhesions with talin1-R8vvv mutant. As fibroblasts and most epithelial cells move with edge protrusion, driven by both actin polymerization and adhesion to the substrate, ERK’s mechanism of control in COS7 cells is expected to be broadly relevant to the many cell types with ERK activation, either through oncogenic mutation or growth factor or substrate environmental inputs (27, 58).

ERK is proposed to function in a protrusion-mediated feedback loop, in which protrusions and associated adhesions trigger ERK activation, which then contributes to protrusion (43, 59, 60). Our data is consistent with this model, in which adhesion engagement leads to paxillin-scaffolded ERK activation, which then phosphorylates adhesion components like paxillin to build the nascent adhesion population. Multiple ERK-dependent phosphorylation sites have been identified on paxillin to be involved in focal adhesion turnover, although which ones are specifically involved in building the nascent adhesion population is unknown (31, 51). Active ERK likely releases from nascent adhesions after substrate phosphorylation and adhesion turnover, diffusing throughout the protrusion and broader cell body. As active ERK builds the short-lived nascent adhesion population and the actin network that pushes against the membrane, the membrane moves forward, and new adhesions form to repeat the process. In this way, adhesions function to amplify ERK activity for cell migration (43, 60). These insights into the mechanisms by which ERK promote protrusion and cell migration form the basis for new understanding in how growth factor and adhesion-activated signaling pathways are integrated to bring about directed cell migration in distinct environments.

## Materials and Methods

### Cell Culture and generation of stable cell lines

COS7 cells were maintained in Dulbecco’s modified Eagle’s medium (DMEM) supplemented with 10% fetal bovine serum (FBS) at 37°C and 5% CO₂. Experiments were conducted on log-phase growth cells, except starvation experiments were in serum-free DMEM for 24 hours. Transfections were performed using TransIT-LT1 (Mirus Bio), at least 48 hours prior to imaging. For migration and imaging assays, cells were plated on acid-washed glass coverslips or glass-bottom dishes (MatTek). AZD6244 (5 µM, Selleckchem) was added to the culture medium 30 minutes prior to imaging. For talin mutant migration and protrusion assays, a reduced concentration of 2 µM AZD6244 was used. EGF (10 ng/mL; Peprotech) was applied to cells 10 minutes prior to fixation or imaging.

The stable EKAR-FAT cell line was generated by co-transfecting COS7 cells with pPB-CMV-EKAR-FAT and HyPBase using TransIT-LT1 (Mirus Bio). Transfected cells were passaged twice and sorted for fluorescence expression with a BD FACS Aria Fusion cell sorter. For paxillin and GIT1 knockouts, COS7 cells were transfected with PXN-1, PXN-3, GIT1-1, or GIT1-2: CRISPR constructs in the pSpCas9(BB)-2A-Puro vector and dilution cloned. Colonies were screened by Western blot, mutations verified by Sanger sequencing, and migration phenotype verified by random migration assay. A luciferase-targeting gRNA was used as a negative control. Representative clones: PXN:1-5 and GIT1:1-4 were used for experiments for further study.

Stable knockdown of talin1 was by short hairpin RNA (shRNA) in pLVX-shRNA1. Lentivirus was produced in HEK293T cells using Lenti-X™ Packaging Single Shots (VSV-G; Takara) and incubated with. COS7 cells for transduction. Cells were selected with 5 µg/mL puromycin and knockdown confirmed by Western blot.

### Plasmids

mCherry-paxillin (#50526), mCherry-talin (#80026), F-tractin-EGFP (#58473), FAT-FAK biosensor (#78303), and mT-Sapphire-C1 (#54545), and pSpCas9(BB)-2A-Puro vector (#62988) were obtained from Addgene. Lentivirus vector pLVX-shRNA1 was from Clontech.

The EKAR-FAT biosensor was derived from pPB EKAREV-NES (EKAREV with a C-terminal nuclear export signal) in Komatsu et al., 2011 and the FAT domain from the FAT-FAK biosensor. The N-terminal focal adhesion targeting (FAT) domain was fused to the N-terminus of EKAREV NES using the EcoRI site with F: AAGAATTCGCCACCATGAGCCTGAACAGCCCCGT GG and R: AAGAATTCGTGGGGCCTGGACTGG.

For bimolecular fluorescence complementation (BiFC) constructs were generated PCR cloning into pcDNA3.1 F/A, using Sapphire from mT-Sapphire-C1 and Venus and Cerulean fragments derived from the FAT-FAK biosensor. For pcDNA3.1-F/A Sapphire VC155-C1-MEK1, the C-terminal fragment of Sapphire was fused to the N-terminus of MEK1 with a GGGSGGGS linker by PCR-cloning of FseI-Sapphire VC155-linker-AscI and AscI-MEK1-NotI. The FseI site and Kozak sequence were added to the Sapphire VC155 fragment with F: AAGGCCGGCCGCCACCATGGACAAGCAGAAGAACGGC and linker and AscI were added with R: AAGGCGCGCCGGATCCACCACCTCCGGATCCACCACCTCCCTTGTACAGCTCGTCCAT GC. MEK1 was PCR-cloned from pcDNA3.1-F/A-MEK1 with AscI and NotI F: AAGGCGCGCCTATGCCCAAGAAGAAGCCGAC and R: AAGCGGCCGCTTAGACGCCAGCAGCATGGG. For pcDNA3.1-F/A Cerulean-N173-N1-PXN, mouse paxillin was PCR-cloned from pcDNA3.1 mApple-paxillin, a gift from the Beckerle lab, and fused to the N173 fragment of Cerulean with the same GGGSGGGS linker. The FseI site and Kozak sequence were added to the paxillin using F: AAGGCCGGCCGCCACCATGGACGACCTCGATGCCC and AscI was added with reverse primer R: AAGGCGCGCCGCAGAAGAGCTTCACGAAGCAG. The AscI site and linker were added to Cerulean N173 with F: AAGGCGCGCCTGGAGGTGGTGGATCCGGAGGTGGTGGATCCATGGTGAGCAAGGGCGG and Not I site added with reverse primer AAGCGGCCGCCTACTCGATGTTGTGGCGGATCTTG. For pcDNA3.1-F/A Venus-N173-N1-GIT1, GIT1 was fused to the N173 fragment of Venus with the GGGSGGGS linker. The FseI site and Kozak sequence were added to the GIT1 using forward primer F: AAGGCCGGCCGCCACCATGTCCCGAAAGGGGCCG and AscI was added with reverse primer R: AAGGCGCGCCCTGCTTCTTCTCTCGGGTG. The AscI site, linker, and NotI sites were added to Venus N173 with the same forward and reverse primers used for Cerulean. The PCR products were cut with FseI, AscI, and NotI and subjected to 3-part ligation with pcDNA3.1 F/A, cut with FseI and NotI.

pcDNA-mNG-Talin1 (shRNA-resistant wild-type talin1 with mNeonGreen) and pcDNA-mNG-Talin1-R8vvv (shRNA-resistant mutant talin1 with the R8vvv substitution) were gifts from Kevin Dean. For lentiviral expression of talin1 shRNA, pLVX-shRNA1-Talin1 was generated by synthesizing complementary oligonucleotides encoding the shRNA described in Han et al., 2021 (IDT, F: GATCCGGAAAGCTTTGGACTACTATTCAAGAGATAGTAGTCCAAAGCTTTCCTTTTTTACGCGT and R: AATTCACGCGTAAAAAAGGAAAGCTTTGGACTACTATCTCTTGAATAGTAGTCCAAAGCTTTCCG) and subjecting the oligos to annealing and ligation into the pLVX-shRNA1 vector using BamHI and EcoRI sites.

For CRISPR knockdowns, guide RNA oligonucleotides targeting PXN or GIT1 were synthesized, annealed, and ligated into the pSpCas9(BB)-2A-Puro vector (PXN-1: GGTCCGGCCATGGACGACCT, PXN-3: CCGTCCAGCGAGGCCCTCAA, GIT1-1: GGCTGGGCATCCATCAGCAG, and GIT1-2: GCACACGCTTGCCAGCAACG).

All constructs were sequence-verified by whole-plasmid sequencing (Plasmidsaurus).

### Antibodies and labels

Primary antibodies were anti-paxillin rabbit monoclonal (Cell Signaling Technology [CST], #2542, 1:1,000), anti-paxillin mouse monoclonal (Invitrogen AHO0492, IF 1:1,000), anti-GIT1 mouse monoclonal (Novus NBP2-22423, clone S39B-8, 1:1,000), anti-GIT1 mouse monoclonal (QED Biosciences #56537, 1:1,000), anti-GIT1 rabbit polyclonal (CST #2919, 1:1,000), anti-talin rabbit polyclonal (CST #4021, 1:500), anti-talin mouse monoclonal (Abcam ab11188, IF 1:1,000), anti-vinculin mouse monoclonal (Abcam ab18058, 1:1,000), anti-GFP mouse monoclonal (Clontech #632381, 1:3,000), anti-ERK mouse monoclonal (Abcam ab54230, 1:500), anti-ERK rabbit polyclonal (CST #9102, 1:1,000), anti–phospho-ERK (Thr202/Tyr204) mouse monoclonal (CST #9201 and #9206, 1:1,000), and anti-GAPDH mouse monoclonal (Ambion AM4300, 1:5,000). Secondary antibodies were Alexa Fluor 568–conjugated goat anti-rabbit (Invitrogen A11011, 1:10,000), Alexa Fluor 647–conjugated goat anti-mouse (Invitrogen A21245, 1:10,000), biotin-conjugated goat anti-rabbit IgG (Invitrogen #31820, 1:1,000), and Alexa Fluor–conjugated streptavidin (Thermo Scientific S11226, 1:500). Nuclei were counterstained with DAPI (Thermo Scientific D1306, 14 µM) or Draq5 (Thermo Fisher 62251, 2 µM).

### Western Blotting and Phos-tag SDS-PAGE

Cells were washed once with cold PBS before lysis and lysed in RIPA buffer supplemented with 0.5 mM EGTA, 1 µg/mL leupeptin, 1 mM DTT, 1 µg/mL pepstatin A, 1 µg/mL aprotinin, 1 mM PMSF, 10 mM sodium pyrophosphate, and 1 mM sodium orthovanadate. For Phos-tag SDS-PAGE, EGTA was omitted from the RIPA buffer. Protein concentration was normalized using a BCA assay (Thermo Scientific) and equal amounts of protein were separated by SDS-PAGE or Phos-tag SDS-PAGE with Mn²⁺-Phos-tag acrylamide (Wako Chemicals). Proteins were transferred to 0.2 µm PVDF membranes, probed with primary antibodies, and detected using IRDye-conjugated secondary antibodies. Blots were imaged using an Odyssey CLx imaging system (LI-COR) and quantified in Fiji/ImageJ.

### Immunofluorescence (IF) and colocalization analyses

For immunofluorescence staining, cells were fixed with 1% formaldehyde (Polysciences) in PHEM buffer (60 mM PIPES, 25 mM HEPES, 10 mM EGTA, 4 mM MgSO₄, pH 6.9), permeabilized with 1% CHAPS in PHEM supplemented with 50 mM β-glycerophosphate and 0.2 mM sodium orthovanadate, and blocked with 20% normal goat serum (NGS, Invitrogen) in MBST (50 mM MOPS, 150 mM NaCl, 0.05% Tween-20, pH 7.4). Primary antibodies were diluted in 5% NGS in MBST as follows: phospho-ERK, rabbit monoclonal (CST #9201) anti-talin (Abcam ab11188), anti-GIT1 (QED Bioscience #56537), and anti-paxillin (Invitrogen AHO0492). Secondary detection for p-ERK was by biotin-conjugated goat anti-rabbit IgG (Invitrogen #31820) and Alexa Fluor–conjugated streptavidin (Thermo Scientific S11226) in 5% NGS in MBST.

For colocalization analysis, COS7 cells expressing mScarlet-paxillin and EKAR-FAT were imaged were imaged on a Nikon Ti-Eclipse microscope with a Nikon Apo TIRF 60× oil objective (NA 1.49), Vortran VersaLase laser launch, Nikon T-FL-TIRF2 TIRF illuminator, Lumencore SOLA light engine, ASI FW-1000 filter wheel, and Photometrics Prime BSI sCMOS camera controlled by MetaMorph software. Consistent filters and acquisition parameters were applied across conditions. Adhesion markers were imaged in TIRF and DAPI by widefield DAPI, with Ex 440nm/ Em 527nm and Ex 561nm/Em 600nm. Pairs of background-subtracted images were analyzed using FIJI JaCoP plugin for Mander’s overlap coefficient using Costes automatic thresholding, the tM1 representing the fraction of the signal from channel 1 overlapping with channel 2 and tM2 representing the fraction of the signal from channel 2 overlapping with channel 1.

### FRET Imaging and analysis

For widefield imaging, cells expressing were imaged on a Nikon Ti-Eclipse inverted microscope with a Nikon Apo TIRF 60× oil immersion objective (NA 1.49), Perfect Focus system, and Photometrics Prime BSI sCMOS camera. Excitation was by Lumencore SOLA light engine. Donor excitation was used a 445 nm CFP filter and emission was recorded sequentially through 480/40 nm or 525/50 nm filters mounted on an ASI FW-1000 filter wheel. TIRF-FRET imaging was performed using the same microscope with either a Nikon T-FL-TIRF2 or Gataca iLas 360 TIRF illuminator. Excitation was by 445 nm laser via a Vortran VersaLaser laser launch. Emission was filtered through a Chroma 91032-Z440/514/561 multiband TIRF cube and passed through either 480/40 nm or 525/50 nm filters on the ASI FW-1000 filter wheel. Imaging exposure, gain, binning, and laser power were constant across channels, using MetaMorph or VisiView software.

FRET image analysis was performed in MATLAB using the Danuser Lab’s Biosensor Analysis Package Version 2.2.0 (https://github.com/DanuserLab). Image stacks were cropped to ROIs. Flat-field and background corrections were performed using signal from empty regions of the coverslip. A binary background mask was generated by thresholding each channel and retaining overlapping regions. Sensitized emission FRET ratios were calculated by dividing the acceptor signal by the donor signal on a per-pixel basis. Photobleaching correction was applied by fitting a single-or double-exponential decay to the average FRET/donor signal ratio across time. Processed images were output as 16-bit TIFFs, with a scale factor of 1,000 applied to the FRET output channel to enable four-digit precision.

### Adhesion Line Scan Analysis

Images were preprocessed in Fiji (ImageJ) to synchronize channels and crop ROIs at the protruding cell edge. Background subtraction, masking, and FRET ratio calculations were performed in the MATLAB Biosensor Analysis Package (https://github.com/DanuserLab). Peak positions of FRET ratios were interpolated by first aligning the FRET ratio.tif and relevant intensity channels in ImageJ, then drawing a line perpendicular to the protruding edge (from cell exterior to interior) through small, assembling adhesions. The fluorescence values across the line were normalized to peak intensity in MATLAB and scans were aligned to the peak of a fiducial marker channel (e.g., mCherry–talin or mCherry–paxillin). Co-localized channel peaks were identified by fitting second-order polynomials to points on either side of the maximum; the vertex of the resulting quadratic curve was taken as the peak position. <u>Migration Assays</u>. COS7 cells were plated and transfected on acid-treated glass-bottom plates. Cells were imaged in FluoroBrite DMEM (Invitrogen) supplemented with 10% FBS and 10 mM HEPES, by phase contrast and widefield fluorescence microscopy using a Nikon Ti inverted microscope equipped with a Plan Fluor ELWD 20× air objective, Lumencore SOLA light engine, Andor Clara CCD camera, and an Oko Labs environmental chamber maintained at 37°C and 5% CO₂. Images were acquired every 10 minutes for 12 hours using Nikon Elements software. For scaffold knockout assays, cells were labeled with Draq5 for automated tracking with MATLAB-based Autocell software (https://github.com/MendozaLabHCI/Autocell). For talin mutant assays, mNeonGreen-expressing cells were manually selected for inclusion after tracking in CellPose (Cyto3 model). Cells exhibiting a persistent random walk were included in the velocity and directionality calculations, with velocity as total displacement over time and directionality the ratio of net displacement to total path length for each time point, averaged over the population. Distributions were compared using the two-sample nonparametric Kolmogorov–Smirnov test.

### Protrusion Analysis

For combined protrusion and adhesion imaging, COS7 cells were co-transfected with mCherry–paxillin and EGFP–F-tractin. Time-lapse TIRF imaging was performed using a Nikon Ti-Eclipse inverted microscope equipped with a Nikon Apo TIRF 60× oil immersion objective (NA 1.49), Perfect Focus system, Nikon T-FL-TIRF2 illuminator, and environmental control for 37°C and 5% CO₂. Fluorescence excitation was provided by a Vortran VersaLase 561 nm laser, and images were acquired with a Photometrics Prime BSI sCMOS camera, controlled by MetaMorph software, at 5-second intervals for 15 minutes. For talin1-knockdown cells, imaging was performed on a Nikon Ti inverted microscope with a Nikon Apo TIRF 60× oil immersion objective (NA 1.49), Yokogawa CSU X-1 spinning disk confocal scanner, and a 488 nm laser. Images acquisitions were at 5-second intervals for 8 minutes using a Photometrics Myo CoolSnap CCD camera with 2×2 binning. Image sequences were analyzed in MATLAB using previously described software (Mendoza et al., 2011; Mendoza et al., 2015). Population distributions were evaluated using the two-sample nonparametric Kolmogorov–Smirnov test.

### Bimolecular Fluorescent Complementation (BiFC)

COS7 cells were co-transfected with BiFC constructs including: pcDNA3.1-F/A BiFC Sapphire VC155-C1-MEK1, pcDNA3.1-F/A BiFC Cerulean-N173-N1-PXN, and pcDNA3.1-F/A BiFC Venus-N173-N1-GIT1. Cells were imaged on a Nikon Ti-Eclipse inverted microscope with a Nikon Apo TIRF 60× oil immersion objective (NA 1.49), Perfect Focus system, Vortran VersaLase 445 nm laser launch, Nikon T-FL-TIRF2 illuminator, and a Photometrics Prime BSI sCMOS camera, with emission filters (480/40 nm, 525/50 nm, and 600/50 nm) mounted on an ASI FW-1000 filter wheel. Imaging Exposure, gain, binning, and illumination intensity were constant across all channels.

Background subtraction was applied prior to analysis. Line scans were drawn perpendicular to the cell edge, bisecting nascent adhesions. Fluorescence intensities were measured for each channel, and peak complementation signal was validated by (1) intensity significantly above background (defined as the mean of 8 pixels, 5 pixels interior from the signal peak) and (2) co-localization with a fiducial marker peak. Background line scans from unrelated cell edge regions were included as controls.

### Adhesion Classification

TIRF imaging of adhesions and adhesion-localized biosensors was performed as described above. Adhesions were classified using a MATLAB-based implementation of the Han lab Focal Adhesion Analysis Package (https://github.com/sangyoonHan (16)). Time-lapse images were cropped to ROIs and sorted by channel. Flat-field correction and background masking were performed and adhesions within 5 µm of the leading edge were analyzed for assembly dynamics and classified based on intensity, lifetime, and edge proximity.

### Graphical Representation Statistical Analysis

Representative images were cropped and linearly adjusted for brightness and contrast using FIJI/ImageJ. Graphs and plots were generated in MATLAB R2023b, GraphPad Prism, and FIJI/ImageJ. Fluorescent images from Western blots and microscopy were acquired using LI-COR Odyssey software (LI-COR Biosciences) or Azure Biosystems Sapphire FL. Figures were assembled in Adobe Illustrator (v29.1). Statistical tests were conducted in GraphPad Prism and MATLAB. Figure legends indicate statistical tests used and the number of replicates. Box plots indicate the median (center line), interquartile range (box), and 95% confidence intervals (notches). Data from two independent groups were compared using a two-tailed unpaired t-test with Welch’s correction when variance was unequal. Paired datasets were first tested for normality and a paired t-test or the Wilcoxon matched-pairs signed rank test was used, as appropriate. For population distributions with undefined variance and for which extreme values may carry biological significance, the Kolmogorov–Smirnov test was used to compare continuous distributions. P-values are reported as p<0.05 (*), p<0.01 (**), p<0.001 (***), and p<0.0001 (****).

## Acknowledgements and Funding Sources

The research reported herein utilized the University of Utah Health Science Center Flow Cytometry, Mutation Generation and Detection, and Cell Imaging Core Facilities. The research was funded by National Institute of General Medical Sciences of the National Institutes of Health through R01GM141372 to MCM and R15GM135806 to SJH. We thank Michiyuki Matsuda, Kyoto University, for providing EKAREV and HyPBase. We thank Kevin Dean, University of Texas, Southwestern for providing shRNA resistant pcDNA-mNG-Talin1 and pcDNA-mNG-Talin1-R8vvv.

## Supplemental Data

**Supplemental Figure 1:**
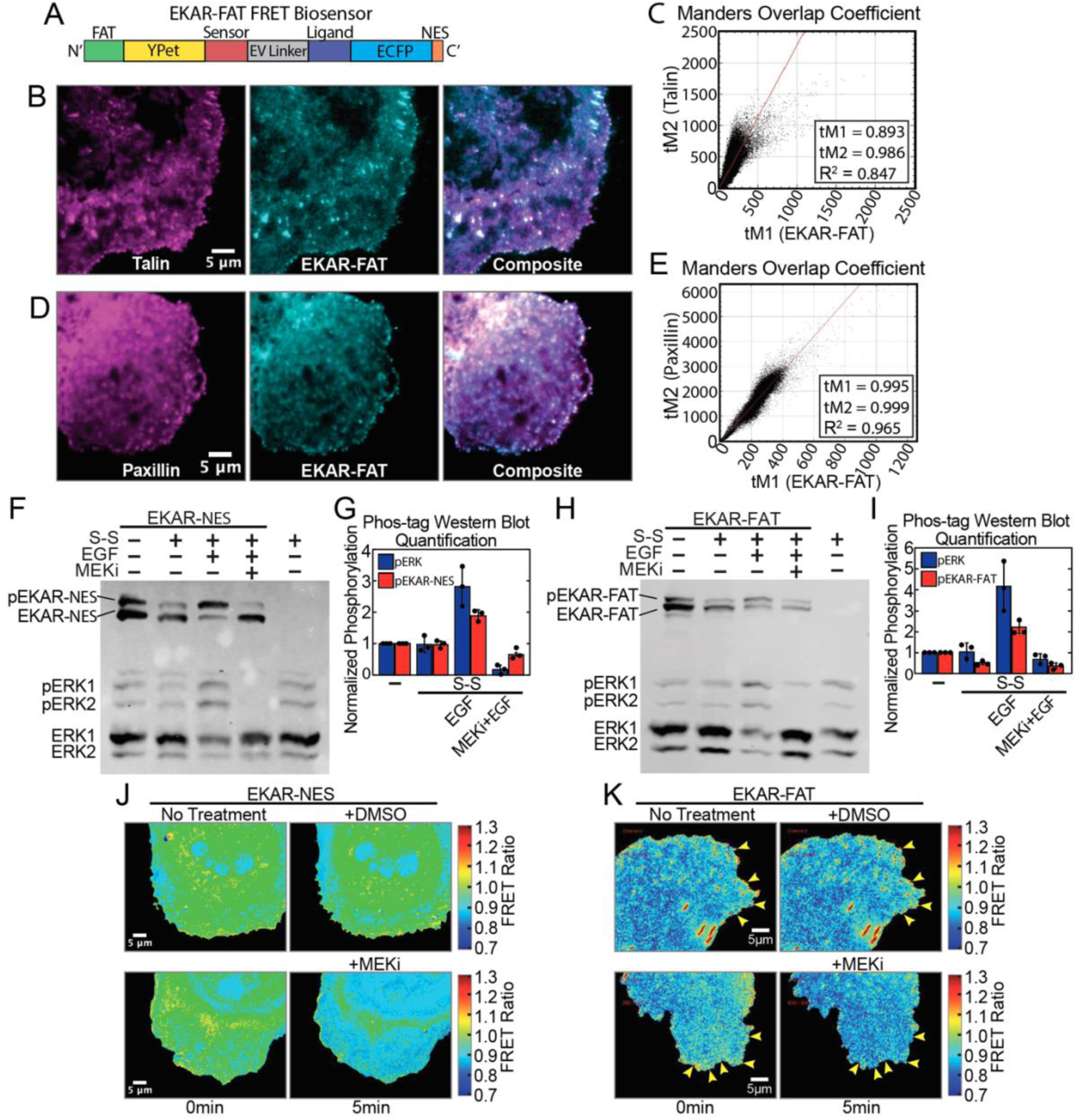
EKAR-FAT senses ERK activity in adhesions (**A**) Schematic of EKAR-FAT FRET biosensor. (**B, C**) TIRF images of COS7 cells transfected with mCherry-talin and EKAR-FAT (CFP emission) and Manders overlap coefficient, calculated from the fluorescent signals from the EKAR-FAT donor (CFP) and mScarlet-paxillin channels. (**D, E**) TIRF images (60x) of COS7 cells transfected with mScarlet-paxillin and EKAR-FAT (CFP emission) and Manders overlap coefficient, calculated from the fluorescent signals from the EKAR-FAT donor (CFP) and mScarlet-paxillin channels. (**F**) Representative Phos-tag Western blot of COS7 cells transfected with EKAR-NES and cultured under log-phase growth conditions or serum-starved and stimulated with EGF for 10 min, with or without MEKi AZD6244. Non-transfected cells control for EKAR-NES identification. (**G**) Phos-tag quantification of D, median values of n=3 experimental replicates. Error bars are SD. (**H**) Representative Phos-tag Western blot of COS7 cells stably expressing EKAR-FAT under log-phase growth conditions or serum-starved and stimulated with EGF for 10 min, with or without MEKi AZD6244, 5 μM. (**I**) Phos-tag quantification of F, median values of n=3 experimental replicates. Error bars are SD. (**J**) Epifluorescence FRET images (60x) of COS7 cells expressing EKAR-NES at before and 5 min post DMSO or MEKi treatment. (**K**) TIRF FRET images (60x) of COS7 expressing EKAR-FAT before and 5 min post 5 µM DMSO or MEKi treatment. FRET ratios are normalized to t=0 min shown in pseudo-color scale. Scale bars are 5 µm.

**Supplemental Figure 2:**
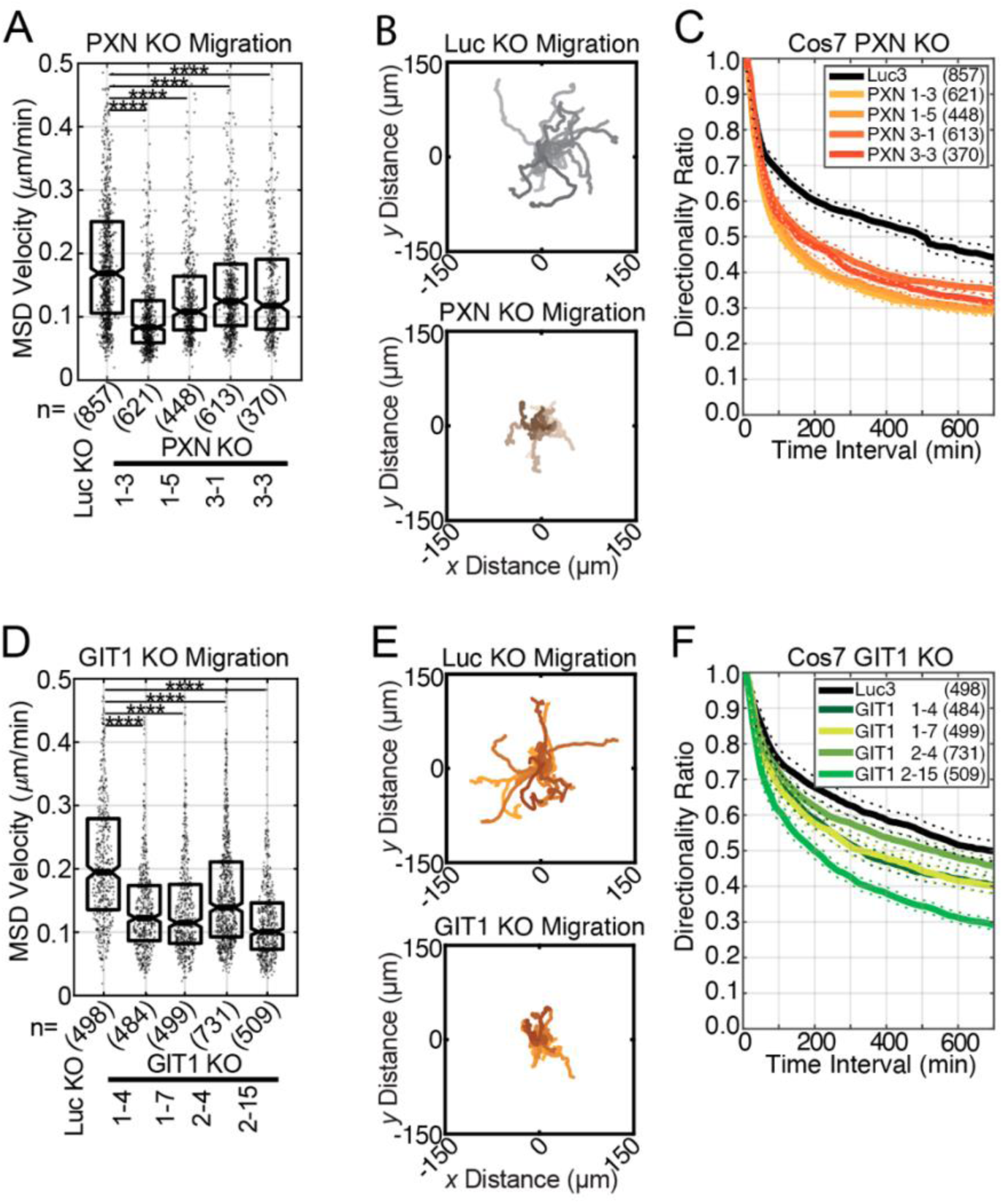
ERK adhesion scaffolds are required for cell migration Random walk migration assay. **(A)** Mean squared displacement (MSD) velocity of control COS7 cells transfected with Crispr targeting Luciferase (Luc KO) cells and four PXN KO clones. Each data point is for an individual a cell, with n cells tracked from three independent experiments. K-S test for significance. Boxes are 25^th^ to 75^th^ distribution. Horizontal lines are medians and notches 95% CI around the median. \*\*\*\**p*<0.0001. **(B)** Rose plots display 20 cell tracks with speeds nearest to the median, Luc KO and PXN KO 1-5. **(C)** Mean directionality of cells in (A), plotted for each time interval. Dashed lines show SEM. **(D-F)** MSD velocity of Luc KO cells and four GIT1 KO clones, rose plots, and directionality as in (A-C).

**Supplemental Figure 3:**
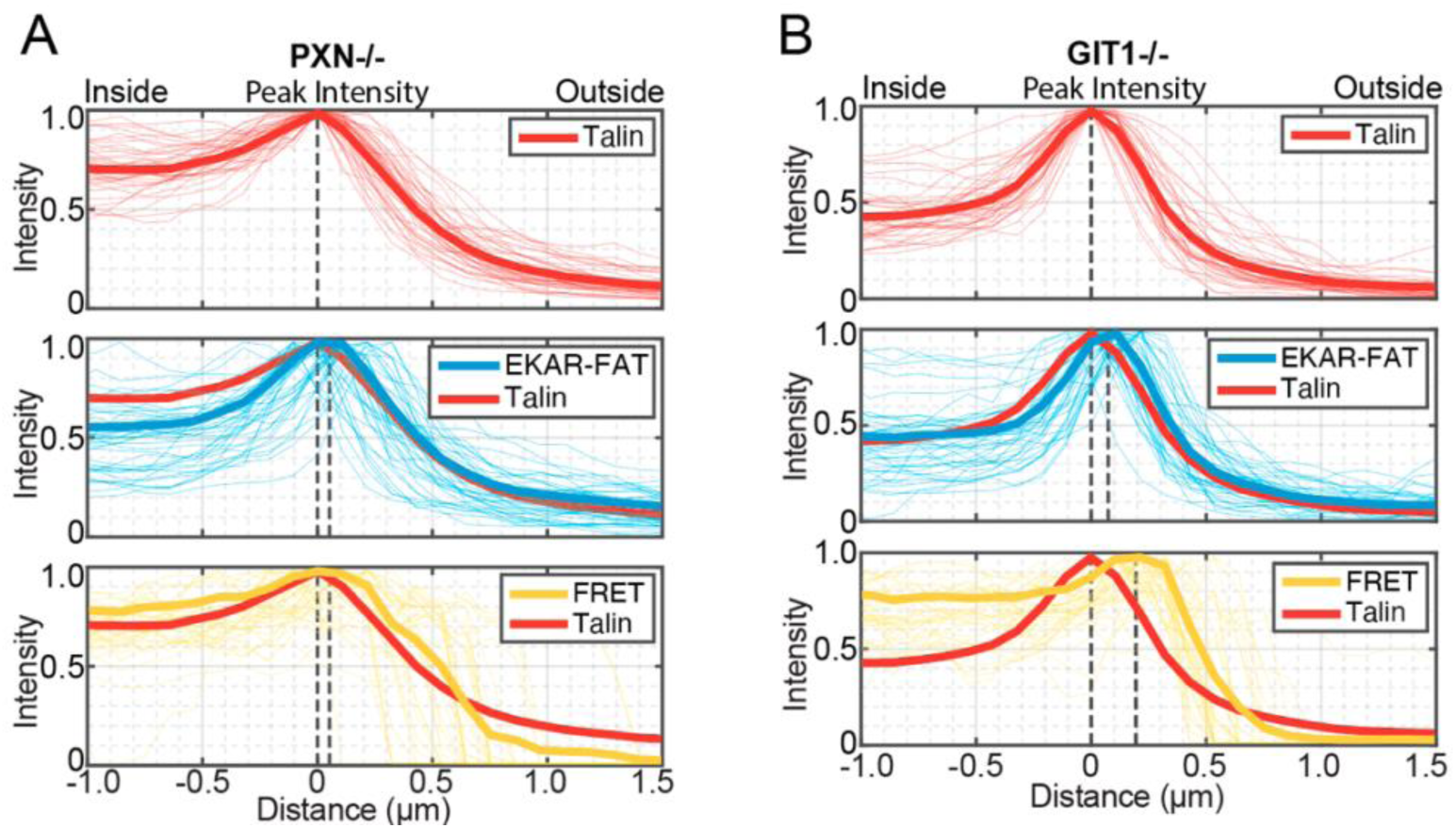
Scaffolds control the localization of Erk activity in nascent adhesions **(A, B)** Normalized peak intensities of line scan data in Fig. 3G, H. Line scans were normalized and aligned in space to the mCherry-talin line’s peak intensity and position, respectively. Peak positions calculated from 3-point polynomial fitting. Bold line is mean. Plots of paired signals of normalized peak intensities from the EKAR-FAT acceptor channel (biosensor localization) and FRET ratio are plotted relative to mCherry-talin (m=40 adhesions from n=5 cells).

**Supplemental Figure 4:**
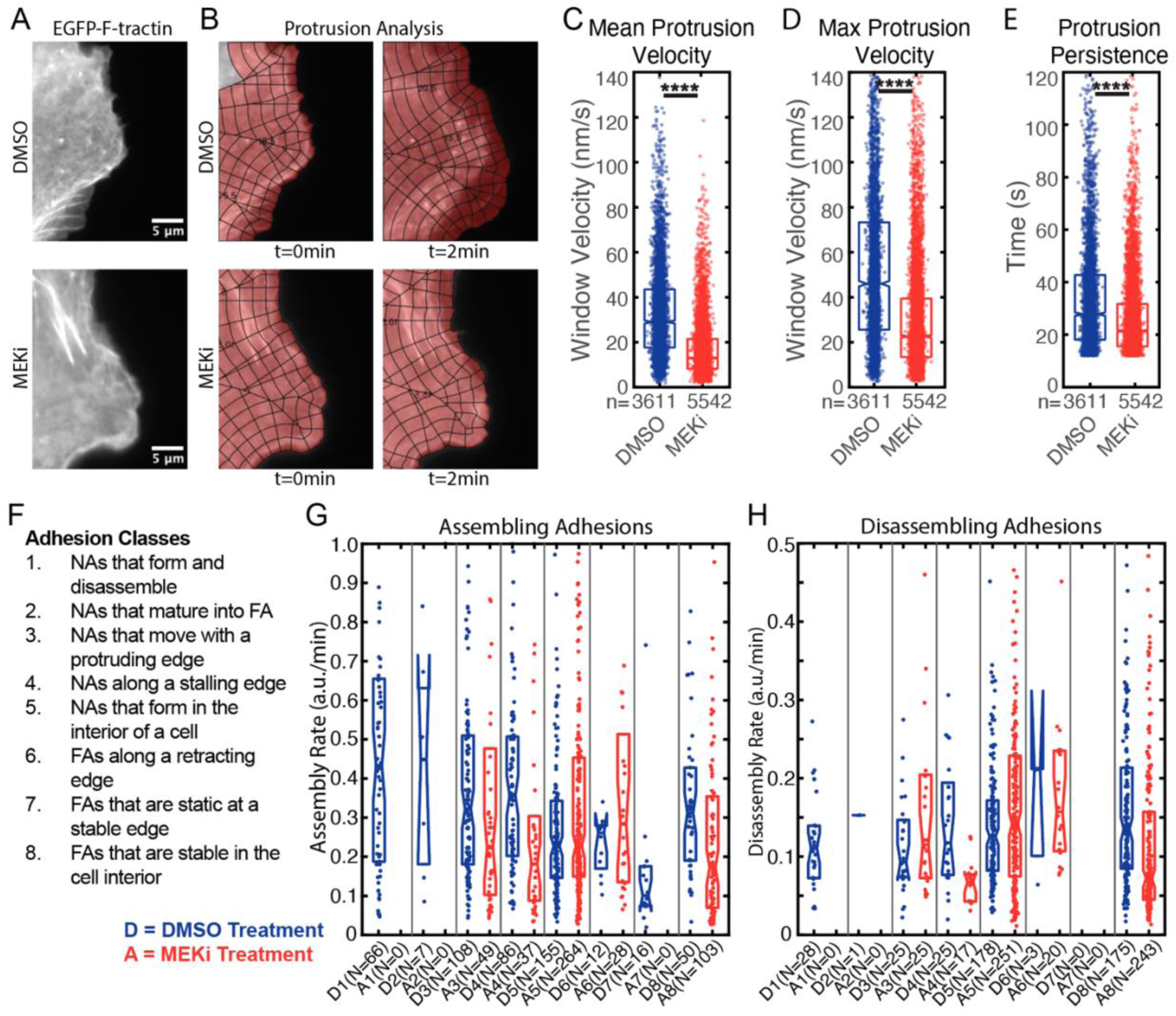
Protrusion and adhesion tracking in COS7 cells treated with MEKi (**A**) Representative images of COS7 cells transfected with EGFP-F-tractin and mCherry-paxillin. Cells were treated with DMSO or MEKi (AZD6244, 5 μM). Movies were acquired for 15 min at 3 s intervals using TIRF microscopy (60x) from n=6 cells from each condition. (**B**) Representative images show of the tracked cell edge from the F-tractin channel in (A), in which ROIs of active leading-edge protrusion were cropped for analysis, segmented, and windowed to provide vector data for edge analysis. (**C**) Mean protrusion velocity, (**D**) maximum protrusion velocity, and (**E**) protrusion persistence. K-S tests. (**F**) Summary of adhesion classes described in Han, *et al.*, 2021. (**G, H**) Extended data set of all classes of assembling and disassembling adhesions from Figure 4. K-S test. Boxes show the 25^th^ to 75^th^ distribution. Horizontal lines are medians and notches indicate 95% CI around the median. \*\*\*\**p*<0.0001.

**Supplemental Figure 5:**
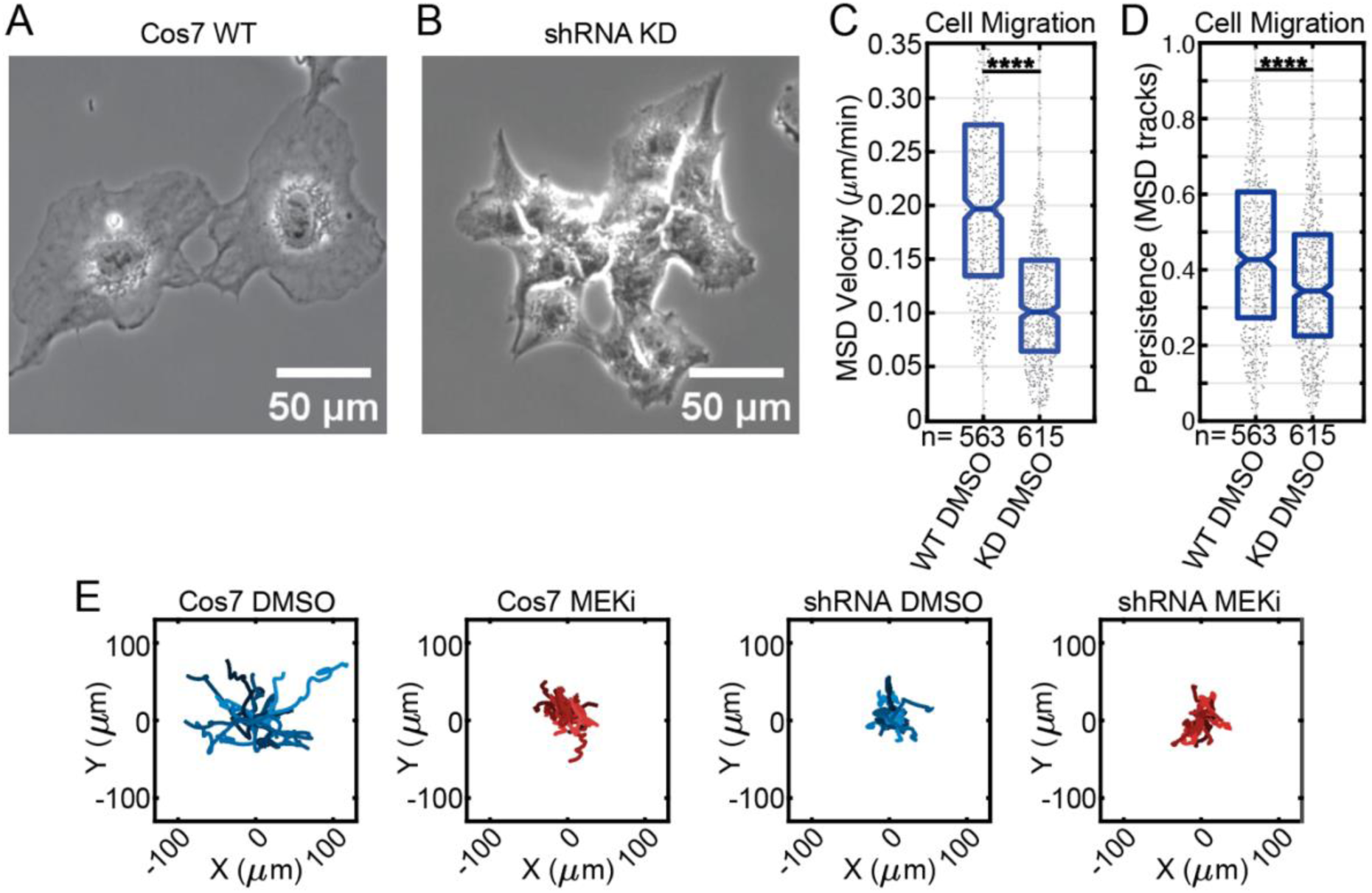
Talin knockdown reduces edge protrusion and cell migration. **(A, B)** Representative phase contrast microscopy images (10x) of cells WT COS7 cells and COS7 Talin1-shRNA KD cells. **(C, D)** Random walk migration assay. Cells were imaged at 10 min intervals for 12 hours (20x phase contrast) and tracked using AutoCell software. K-S test. Boxes show 25^th^ to 75^th^ distribution. Horizontal lines are median, and notches show 95% CI around the median. \**p*<0.05, \*\**p*<0.01, \*\*\**p*<0.001, \*\*\*\**p*<0.0001. **(E)** Rose plots display 20 cell tracks with speeds nearest to the median for each condition.

Supplemental Movie 1 - DMSO Adhesion Classification

Supplemental Movie 2 - MEKi Adhesion Classification

Supplemental Movie 3 - WT Talin MEKi Migration

Supplemental Movie 4 - R8vvv MEKi Migration

